# A Parcellation Scheme of Mouse Isocortex Based on Reversals in Connectivity Gradients

**DOI:** 10.1101/2022.08.30.505842

**Authors:** Michael W Reimann, Timothé Guyonnet-Hencke

## Abstract

The brain is composed of several anatomically clearly separated structures. This parcellation is often extended into the isocortex, based on anatomical, physiological or functional differences. Here, we derive a parcellation scheme based purely on the spatial structure of long-range synaptic connections within the cortex. To that end, we analyzed a publicly available dataset of average mouse brain connectivity, and split the isocortex into disjunct regions. Instead of clustering connectivity based on modularity, our scheme is inspired by methods that split sensory cortices into subregions where gradients of neuronal response properties, such as the location of the receptive field, reverse. We calculated comparable gradients from voxelized brain connectivity data and automatically detected reversals in them. This approach better respects the known presence of functional gradients within brain regions than clustering-based approaches. Placing borders at the reversals resulted in a parcellation into 41 subregions that differs significantly from an established scheme in nonrandom ways, but is comparable in terms of the modularity of connectivity between regions. It reveals unexpected trends of connectivity, such as a tripartite split of somatomotor regions along an anterior to posterior gradient. The method can be readily adapted to other organisms and data sources, such as human functional connectivity.

## 1 Introduction

Established parcellation schemes of the cortex (Zilles and Amunts, 2010; Amunts and Zilles, 2015; Sporns, 2015) use anatomical differences - such as presence of a layer 4 - or functional differences - such as responding to certain modalities with low delay - to draw borders between regions. Connections from other brain structures - such as innervation from different thalamic nuclei - also play a role, but connectivity within cortex is typically analyzed with respect to a prior parcellation scheme (Scannell et al., 1995; Bota et al., 2015). Implicit in this is the assumption that the prior scheme is useful in revealing the structure of connectivity. This assumption is not unwarranted, as the observed functional differences are at least partially the result of the intracortical connectivity, in the sense that differences in functional behavior of neurons across regions is likely to be mirrored by differences in connectivity (van den Heuvel et al., 2015). However, the details of parcellation schemes can quantitatively affect the results and consequently make comparison across studies difficult (de Reus and van den Heuvel, 2013). Therefore, this logic has been reversed, splitting the cortex into regions based on its connectivity (Tittgemeyer et al., 2018; Eickhoff et al.,2015). Then, function or anatomy can be analyzed with respect to the results.

Here, we describe our attempt at connectivity-based parcellation using the average mouse brain connectome published by the Allen Institute (Oh et al. (2014); Knox et al. (2019), connectivity.brain-map.org, github.com/AllenInstitute). It is at this point the most comprehensive source of information on structural mouse brain connectivity. Furthermore, we have previously demonstrated that it is of sufficient quality to describe the spatial structure of connectivity in terms of pathway strengths, layer profiles and topographical mapping, with respect to a prior brain parcellation (Reimann et al., 2019). Nevertheless, our results must be interpreted with the caveat that they may be affected by inaccuracies in this source material - the output can only be as accurate as the input. To that end, we ensure that our algorithms are general enough to be readily applied to future, improved data sources. This includes data sources for different organisms, such as human functional connectivity.

A wealth of clustering algorithms exist that could use the connectivity data as an input and provide a parcellation that optimizes a measure such as similarity of connectivity within a region. Such an approach was typically used in previous publications on connectivity based parcellation (Craddock et al., 2012; Eickhoff et al., 2015; Tittgemeyer et al., 2018). Here, however, we developed an alternative method inspired by existing data analysis techniques that find borders between regions in sensory cortices. These techniques consider gradients of the locations of receptive fields associated with cortical locations and draw borders between regions where sudden reversals of gradients are observed (Sereno et al., 1994; Garrett et al., 2014; Schönwiesner et al., 2015; Juavinett et al., 2017). To find analogous reversals in connectivity data, we used a technique called diffusion embedding to detect the strongest gradients in the similarity structure of the voxelized connectome (Coifman and Lafon, 2006). We found that similar gradient reversals were present in that data that could be automatically detected to determine borders between regions. Eickhoff et al. (2015) pointed out that cortical regions are often functionally organized around gradients (see also Wandell et al. (2007); Hensel et al. (2015)) and that similarity clustering tends to cut borders into them. Our approach avoids this potential issue as it places borders at reversals instead of continuous sections of gradients.

Compared to methods around reversals of functional gradients, the advantage of our approach is that whole brain, or at least whole cortex connectomes are readily available, enabling the generation of a coherent parcellation of the entire system. To achieve the same based on functional data, first the modalities of suitable gradients would have to be identified for each part of the cortex, and then individually probed. Conversely, diffusion embedding of connectivity has been used to separate parts of the mouse brain before, even on the same data set. For example, (Coletta et al., 2020) showed that regions associated with vision, audition, somatosensation, motor activity and the default mode network clearly separate based on the two strongest extracted components. However, more finegrained separation was not shown. Conceptually, our approach is differs drastically in that it specifically considers reversals in the spatial gradients of the embedding results. This affects the output, as even voxels associated with very similar numerical values can be placed in different sub-regions, if they fall on opposite sides of a reversal.

### 1.1 Motivation and approach

Established techniques draw borders between sensory cortical regions by considering the activity of neurons at various locations, specifically the properties of their preferred stimuli (Sereno et al., 1994; Garrett et al., 2014; Schönwiesner et al., 2015; Juavinett et al., 2017). For some properties, this comes in the form of a continuous topographical mapping, meaning that the entire spectrum of values of the property can be associated with specific cortical locations, with nearby locations being associated with similar values. A property can be represented this way several times in neighboring cortical regions, such as sound frequency being represented in auditory cortices (Schönwiesner et al., 2015) or the visual field (retinotopy) being mapped to primary and higher-order visual cortices (Sereno et al. (1994); Garrett et al. (2014); Juavinett et al. (2017); Fig. 1A). At the border between these regions, we cross from one continuous map to another, yet there is no sudden jump in the value of the property, but rather a gradient reversal (Fig. 1B, black arrows). This means that the maps of neighboring regions associated with the same modality are mirror images of each other, which can be used to detect borders between regions. Note that Garrett et al. (2014); Juavinett et al. (2017) specifically use a change of the sign of the gradient of a two-dimensional mapping (horizontal and vertical retinotopy) to detect borders (Fig. 1B, red and blue outlines), but most of the time this change arises from a reversal of one gradient while the other remains constant.

**Figure 1:**
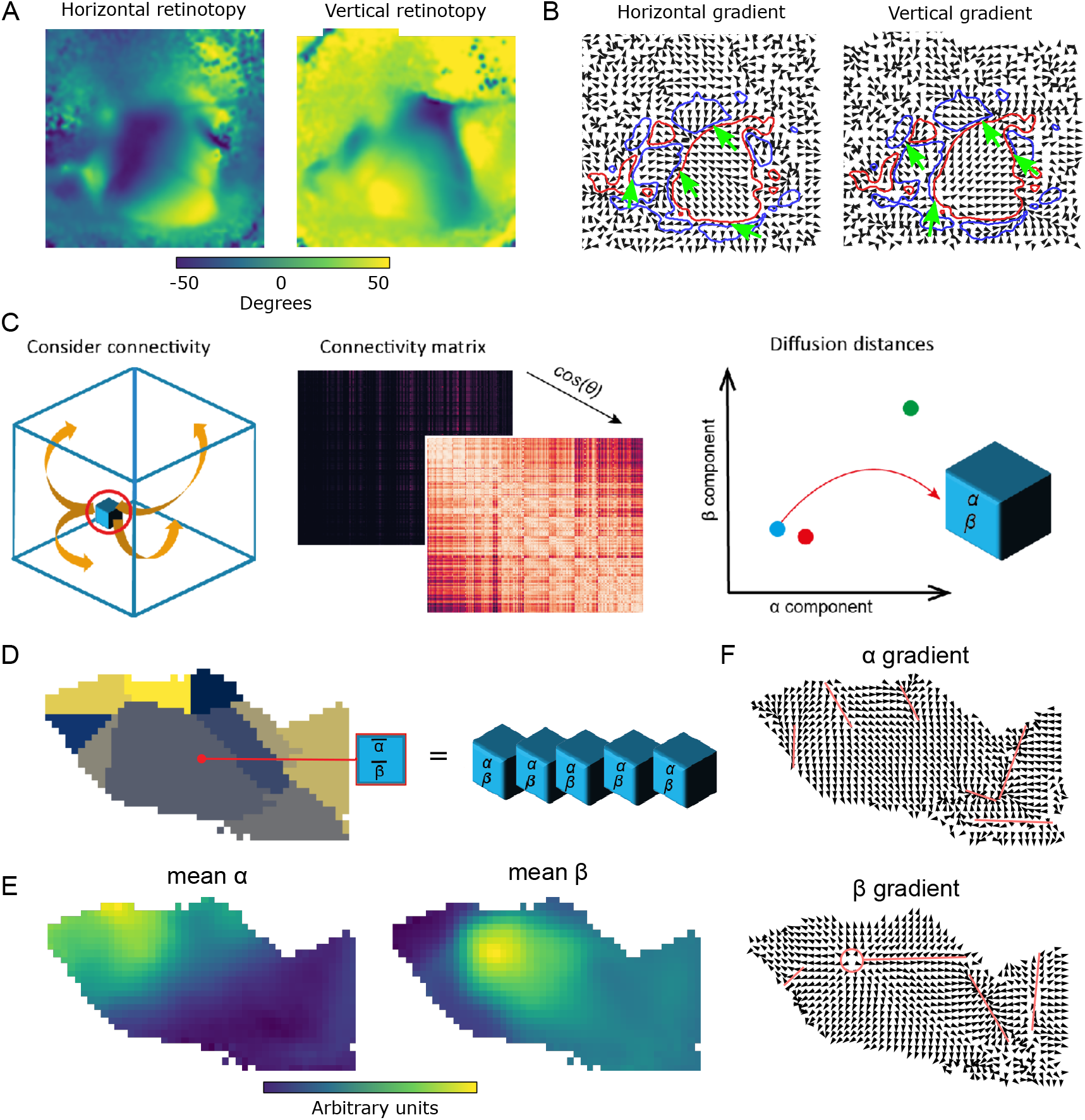
Parcellation at gradient reversals. A: Horizontal and vertical retinotopy of mouse visual regions as measured by Garrett et al. (2014). Data digitized from the original publication. B: Gradients of the retinotopies (black arrows). Overlaid are regions with positive sign (smallest rotation from horizontal to vertical gradient is clockwise, red) and with negative sign (smallest rotation is counter-clockwise, blue). Green arrows denote examples of switches of the sign by a sudden reversal of one gradient. C: Diffusion embedding. Build a similarity matrix based on the connectivity strength of considered voxels and run diffusion process on this matrix to extract the embedded geometrical space of connectivity. Extract then the first two dominant eigenvectors of the diffusion distance indicating the connectivity ‘profile’ of the voxels. D: Left: Flattened view of mouse visual cortex. Subregions according to AIBS CCF indicated in different colors. Right: For each pixel of the flat view we average the first two diffusion coordinates (*α* and *β*) of the voxels mapping into the pixel. E: Resulting mean *α* and *β*. F: Gradients of the mean *α* (top) and *β* (bottom). Red lines indicate manually annotated borders draw at gradient reversals.

We hypothesize that even in the absence of activity recordings, the gradient reversal and thus the border between regions can be detected when considering the connectivity of neurons instead. This is based on the idea that it is the afferent connectivity of neurons, i.e. their input, that determines the quality of their preferred stimulus. In addition, neurons with similar preferences have been demonstrated to connect strongly to each other (Rossi et al.,2020; Harris and Mrsic-Flogel, 2013), indicating that also their targets of synaptic outputs are similar. Thus, we attempted to detect gradients in the profiles of afferent and efferent connectivity of individual voxels in the Allen Institute’s voxelized mouse connectome.

A technique to find such gradients, diffusion embedding (Vos de Wael et al., 2020; Coifman and Lafon, 2006), has been used on this dataset before to characterize it (Coletta et al., 2020). Briefly, it positions every voxel in an n-dimensional coordinate system, such that voxels with similar connectivity profiles (i.e. receiving inputs from the same locations and sending outputs to the same locations) are placed closer to each other (Fig. 1C; for details, see Methods, Fig. S1 and (Vos de Wael et al., 2020)). This process reveals the embedded geometric structure of connectivity within the data. Each dimension of this coordinate system corresponds to a *component* of connectivity with components placed in order of their strength. In that regard, it can be thought of as similar to principal component analysis.

Previous research using this technique indicates that the difference between sensory modalities forms one strong gradient in the dataset (Coletta et al., 2020). This indicates that the principle of gradient reversal might be useful not merely to distinguish sensory regions of the same modality, but also different modalities and even sensory regions from non-sensory associative areas.

## 2 Results

### 2.1 Gradient reversals in visual regions

To test our hypothetical approach, we first investigated whether the well-established gradient reversals in visual cortices can be recreated from the voxelized connectivity. To that end, we gave the output of the diffusion embedding a spatial context that could be used to determine its gradients. We used a *flat map* of mouse cortical regions that associates each connectivity voxel with a discrete location in a two-dimensional grid. The projection preserves the relative area of the surface defined by the layer 4 to layer 5 boundary. Voxels along an axis orthogonal to cortical layers were mapped to the same location that we refer to as a *pixel*. At a voxel resolution of 100μm, 13 ± 8 (mean ±std) voxels were mapped to the same pixel. For each pixel, we considered the mean values of the diffusion embedding coordinates of the voxels mapped to that pixel (Fig. 1D). We could then calculate the gradient of any diffusion embedding coordinate along the x- and y-axis of the pixelized image of isocortex (Fig. 1E, F).

This approach explicitly “flattened away” the cortical layers, and it is known that the laminar structure decidedly shapes connectivity (Felleman and Van Essen, 1991). However, we were interested in a parcellation orthogonal to layer boundaries, similar to existing parcellations and in accordance with the theory of the cortical column, i.e. a piece of cortex spanning all layers acting as a functional unit. We characterized the amount of information lost through averaging by comparing the spread of diffusion embedding coordinates of voxels mapped to the same pixel to the one over the mean values of pixels in the same region (Fig. S2). We found that the standard deviation of values within a pixel was almost two orders of magnitude below the standard deviation over a region, demonstrating that the connectivity profiles of voxels along a cortical column are likely similar, thus avoiding too much information through flattening. Yet, in the future our algorithms might be extended by additionally considering a third gradient of a diffusion embedding coordinate along the depth-axis.

We applied this approach to the voxels associated with visual areas in the Allen Institute Common Coordinate Framework (AIBS CCF, Wang et al. (2020), atlas.brain-map.org, Fig. 1D-F). We found in the two strongest components (from here on called *α* and *β*) several gradient reversals that roughly coincide with the established parcellation. In the *α* component we found reversals at the border between VISam and VISpm, splitting VISrl from VISa and VISal, and a number of reversals splitting VISal, VISl and VISli from the rest. In the *β* component we found a local maximum, leading to a reversal when crossed in any direction, at the point where VISp, VISpm, VISam, VISa and VISrl meet. Additionally, reversals split VISa, VISrl and VISal from the rest. Crucially, the large area of the primary visual cortex (VISp) was not split up by reversals, instead depicting a *continuous mapping* characterized by only smooth changes in the direction of both the *α* and *β* gradients.

We conclude that the diffusion embedding coordinates of voxelized connectivity contain similar gradients and gradient reversals as the functional data based on neuron activity. As such, it may be possible to build a parcellation based on it.

### 2.2 Context-dependence of connectivity gradients

The diffusion embedding process yields connectivity trends ordered by their strengths. This means that the results are potentially dependent on the context, i.e. on which parts of cortex were subjected to diffusion embedding together. A large-scale connectivity trend, spanning the entire isocortex, such as the one described by component *α* in Fig. S3A, will not depict its full range of values when evaluated over a small patch. Consequently it will appear weaker and may even be lost to noise. Conversely, a trend with a higher spatial frequency would be less affected. This indicates that also reversals may appear or disappear depending on the spatial context and scale.

We evaluated the degree of context-dependence by considering the gradients of the two strongest components in an exemplary region (SSp-ll, the lower limb region of primary somatosensory cortex) in different contexts: As part of the entire isocortex (Fig. S3A), the somatosensory and motor areas (same figure, panel B), somatosensory areas (panel C) and individually (D). Note that the values of *α* and *β* are in arbitrary units and consequently their sign is arbitrary as well. This means that only the orientations of gradients and locations of reversals are meaningful, but not their direction. As such, results from the level of isocortex down to somatosensory areas are very similar, only once SSp-ll is considered in isolation does *β* form a second, orthogonal gradient.

The emergence of an orthogonal gradient can be intuitively understood as follows: With the region considered in isolation, we can assume that gradients related to region differences will appear weak or not at all. As such, the strongest gradients would be related to the local connectivity. Its known distance dependence (Petersen and Sakmann, 2000; Lubke, 2003; Holmgren et al., 2003; Perin et al., 2011) will then result in gradients that simply reflect the spatial locations of the voxel. In the flat view, this would result in two continuous (i.e. non-reversing) and orthogonal gradients, as observed.

We conclude, that the orthogonality and continuity of the top two gradients, evaluated in a region in isolation, may be an indicator that the region is *atomic*, i.e. does not require further subdivision. Later, we will further test this hypothesis in a model with a known set of atomic regions. We defined two metrics that measure this trend: *Gradient deviation, gd*, evaluates how much the angle between the two strongest gradients deviates from 90 degrees, and *reversal index, ri*, evaluates the absence of reversals by counting the pairs of pixels with an angle above 90 degrees in the same component (for details, see Methods).

For SSp-ll we found that *gd* decreases significantly over scales (Fig. S3E), as expected. While ri is actually lowest at the scale of somatosensory areas and increases slightly for SSp-ll in isolation, the reduction relative to the entire isocortex is still substantial. Going forward, we will use these metrics to evaluate how distant a proposed parcellation is from the theoretically derived ideal.

### 2.3 Evaluation of an algorithmic detection of reversals

We implemented and executed two approaches to automatically split regions based on connectivity gradients: *cosine distance clustering* and *reversal detection* (Fig. 2A). The first generalizes the process of Campello et al. (2013) by considering the angle between a pair of gradients, then clustering them based on similarity. The second one explicitly determines pixels of the flattened view where a gradient reverses (*border pixels*), then calculates for each pair of pixels the number of border pixels separating them and clusters the resulting distance matrix (for details, see Methods).

**Figure 2:**
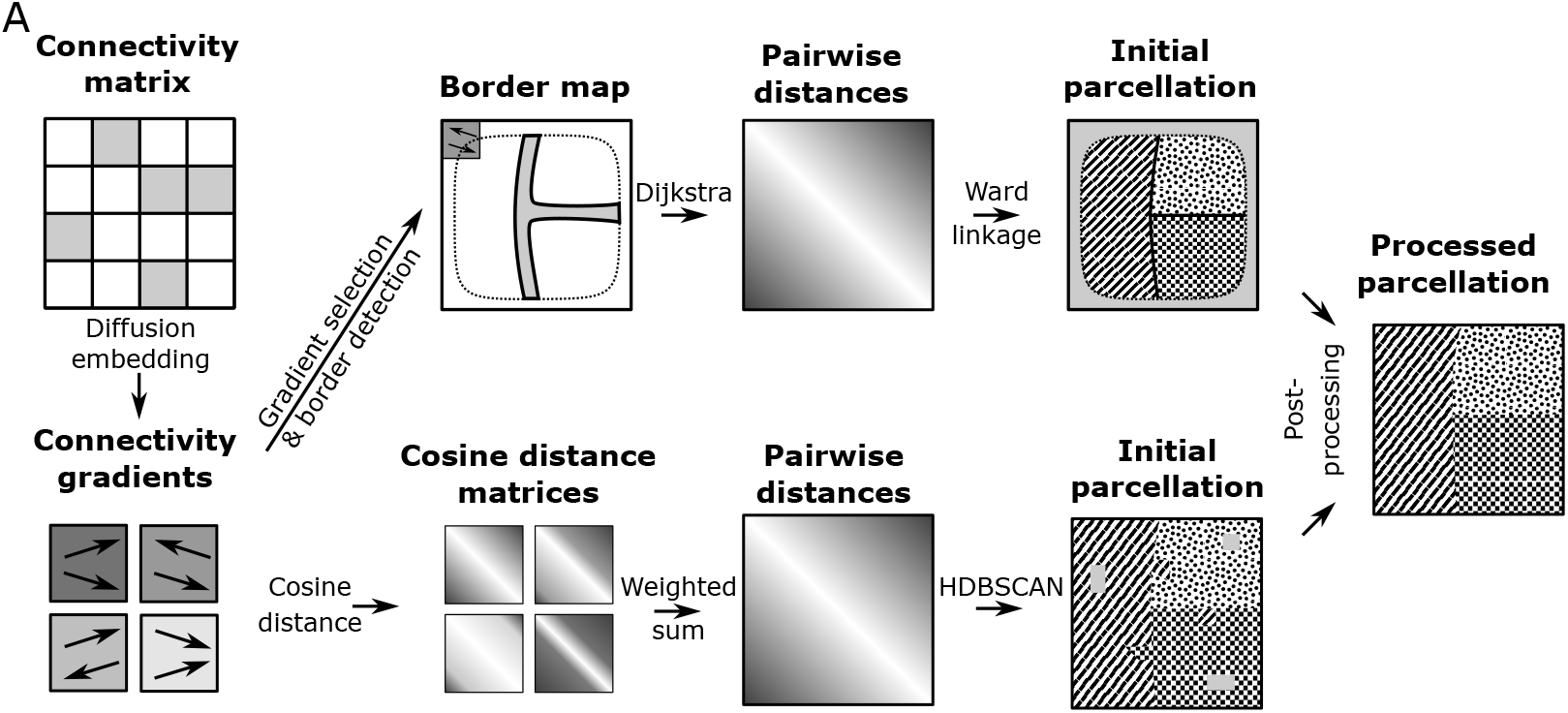
Parcellation methods and evaluation in a toy model of connectivity. Symbolic illustration of the workflow. Input is the matrix of connection strengths between voxels of the region to parcellate. The 20 strongest components are extracted through diffusion embedding and gradients are calculated in a flattened view (four gradients indicated). An initial parcellation is derived with one of two methods. Top: A gradient is selected, based on minimizing the reversal index of the resulting parcellation, and a border map is built by convolving the normalized gradient with a uniform kernel. Pixels where the length of the convolved gradient is below 0.97 are classified as border. Distance between a pair of pixels is calculated as the number of border pixels on the path between them using the Dijkstra algorithm. An initial parcellation is generated through Ward linkage. Bottom: For each gradient pairwise cosine distances of the direction of the gradient are calculated. They are summed with weights corresponding to the strength of the respective gradient. An initial parcellation is generated by the HDBSCAN algorithm. Finally, the initial parcellation is post-processed.

Both approaches had their advantages and disadvantages. *Reversal detection* considered only one component at a time, requiring the selection heuristic. Also, its convolution step left a number of pixels at the borders of the region unlabelled (Fig. 2, top right). *Cosine distance clustering*, while considering several components together, did not explicitly seek out reversals, but rather considered absolute differences in gradients. Results also tended to be noisier, and it potentially left a number of pixels classified as outliers and thus unlabelled (Fig. 2, bottom right). These issues were addressed in a post-processing step that generalized region assignment to unlabelled pixels and ensured that the resulting regions were spatially continuous (see Fig. S6; Methods).

As gradients and their reversals were context-dependent (see above), we did not stop after a single application, but repeated the process recursively on the resulting sub-regions. This would result in additional subdivisions, if new reversals appear at the reduced spatial scale. Further repetitions thereby lead not only to a more fine-grained parcellation, but also a proposed hierarchy of regions.

We evaluated this workflow against toy models of connectivity with a known, ground truth parcellation. The purpose was to better understand how robustly the procedure works on connectivity data with various, interacting gradients. The models were built based on strong assumptions about the organization of connectivity gradients. We do not claim that they reflect biology, only that they are useful for evaluating the ability of our algorithms to detect known gradients and their resistance against noise. We began with two very simplified models, *reversing hierarchy* and *node distance*. Both hierarchically split the unit cube along orthogonal axes into equally sized sub-regions, explicitly assign target gradients that reverse at the borders, and assign connection strengths reflecting the gradient (Fig. S4A). They differed in the way the prescribed region hierarchy was taken into account (Fig. S4B1 vs. B2; for details, see Methods).

We attempted to recreate the parcellations of toy models with three hierarchy levels (Fig. 3A) and white noise at low (*ϕ* = 0.1) or high (*ϕ* = 5.0) amplitudes added to each weight (Fig. S4C; noise amplitude specified relative to the strength of the signal). For the *reversing hierarchy* model at low noise, two successive applications of *cosine distance clustering* or four applications of *reversal detection* recreated most of the parcellation, although not at full granularity in the case of cosine distance clustering (Fig. 3B1, Fig. S5 for results of intermediate steps). Under high noise conditions, only the parcellation up to the second hierarchy level was recovered.

**Figure 3:**
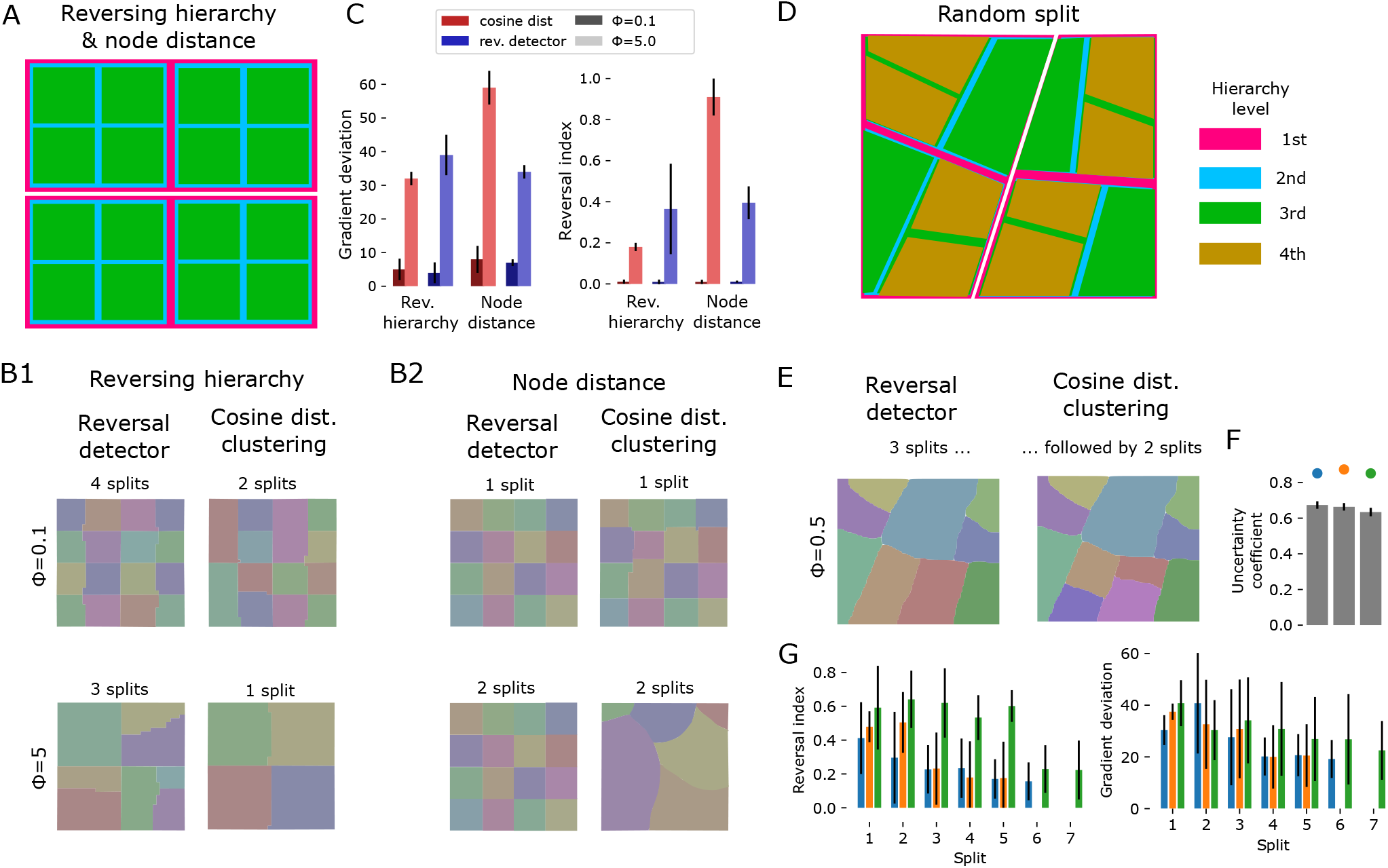
Applying splitting methods to toy models with known ground truth. Toy models of connectivity were constructed as in Fig. 2. A: Reversing hierarchy and node distance models split the unit cube into equal quadrants. B1, left: Results of splitting the reversing hierarchy model using reversal detection, with noise amplitudes *ϕ* = 0.1 (top) and *ϕ* = 5 (bottom). The number of applications of the splitting algorithm before the solution converges is indicated above each result. Right: splitting with cosine distance clustering. B2: Same, for the node distance model. C: Gradient deviation and reversal index of the final output for the different approaches. Shades of red: Using cosine distance clustering at low or high noise levels. Shades of blue: Using reversal detection. Bars and error bars indicate mean and standard deviation over detected regions. D: The random split model splits the volume at randomly drawn lines, connecting points on opposite sides with each other. Exemplary random instance indicated. E: Results for the exemplary instance in D. Left: After three splits with reversal detection. Right: After following reversal detection with two splits using cosine distance clustering. F: Uncertainty coefficient (uc) of the true parcellation and the solution reached by the algorithms for three toy model instances. The instance in E is indicated in orange. Grey bars and error bars indicate mean uc and its standard deviation for random control parcellations with the same numbers of regions. G: Reversal index (left) and gradient deviation (right) after successive splits for three instances.

Conversely, for the *node distance* model (Fig. 2C2), the full parcellation was reconstructed after a single split in the low noise condition (Fig. 3B2). Under the high noise condition, reversal detection required two applications, and the output of cosine distance clustering did not reflect the true parcellation in any way.

We conclude that our methods are likely to recreate at least part of a parcellation, and that resistance of the method against noise depends on the underlying architecture of connectivity, which still remains unknown. Further, the ways in which the algorithm can fail is by yielding an incomplete parcellation, or a completely incorrect parcellation. However, note that the more severe, second mode of failure was detectable through the quality metrics. Both *ri* and *gd* of the results were higher for the incorrect solution than for the correct ones. Especially *ri* appeared to be a good evaluator, yielding a value over two times (0.87 vs 0.39) higher in the case of the incorrect solution (Fig. 3C). For the combination of reversal detection in the reversing hierarchy model under high noise conditions, the recovered parcellation was incomplete in some parts and complete in others; this was reflected by a large standard deviation of values of *ri*.

We evaluated to what degree the above generalizes to less idealized conditions in a more complex model that generates random, more irregularly shaped sub-regions with different sizes (Fig. S4D, see Methods). We evaluated it at an intermediate noise level of *ϕ* = 0.5 (Fig. S4E). Based on our experiences with the simpler models, we began with applications of the more conservative reversal detection until it recovered no further granularity (Fig. 3E, left), followed by cosine distance clustering for further granularity (Fig. 3E, right). Values of the reversal index and gradient deviation were lowered at each split to around 0.2 and 20 degrees respectively (Fig. 3G). Similarity of the results and the true parcellation, measured by their uncertainty coefficient was high (Fig. 3F), yet the results were imperfect in two ways: Some borders were slightly shifted, and some separate sub-regions were merged into one.

### 2.4 Applying the algorithms to mouse cortical connectivity

We finally applied the splitting algorithm to the connectivity of the entire mouse isocortex. In all of the six strongest components, except for the first and third, a prominent reversal was immediately visible, spanning horizontally across the flat view (Fig 4A). We applied a first split using *reversal detection*, as its results appeared more robust during our evaluations. The method identified the sixth component as the one to use to minimize the result ri. This component had 37% of the strengths of the first, strongest component (Fig. 4B).

**Figure 4:**
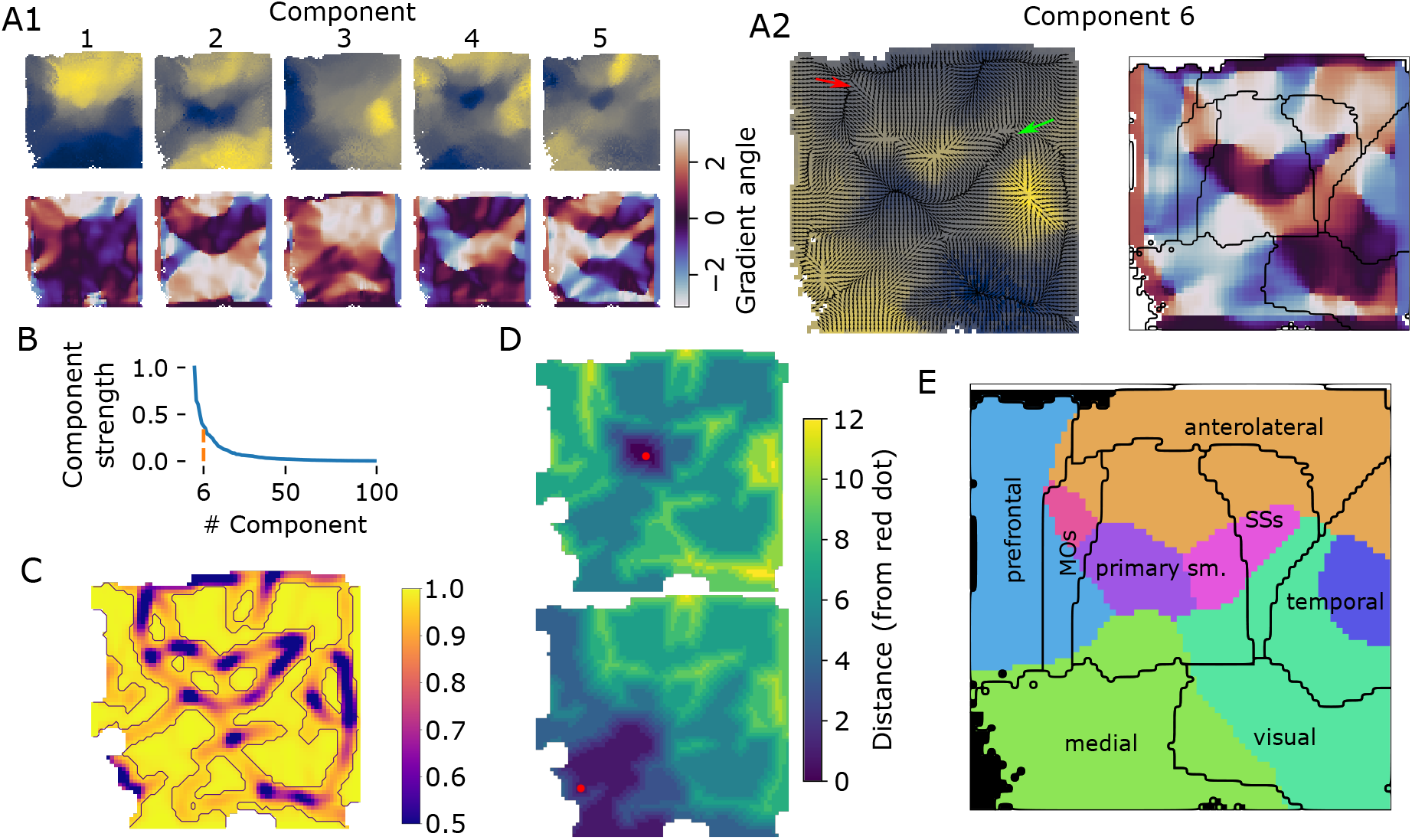
First split of isocortex into subregions. A1: Top: Raw values of the first five diffusion component of mouse isocortex. Bottom: Angle of the gradient at each location. A2: Left/right as A1 top/bottom for the sixth component that was selected for the split. Overlaid in black, left: Gradient field, with each vector normalized to unit length; right: Established rough parcellation (modules of Harris et al. (2019), but with MOs and SSs separated) B: Strengths of components relative to the strongest. Orange: the selected sixth components has a strength 37% of the top one. C: Lengths of vectors resulting from convolving the field in A2 with a two-dimensional Gaussian kernel with a width of five pixels. Purple outlines: Pixels with a vector length < 0.97, thus classified as a potential border. D: Resulting distance metric; drawn is the distance between any pixel and for two exemplary pixels indicated by red dots. E: Resulting first split (colored areas), with established parcellation overlaid and labelled.

Closer inspection of the reversals of that component (Fig. 4A2) and comparing to an established parcellation, revealed several strong reversals. This included one starting between the prefrontal and anterolateral regions (Fig. 4A2, red arrow), cutting through somatomotor regions, and ending at the point where anterolateral, somatomotor and temporal regions meet (green arrow). This led to borders being detected that surround compact regions within the somatomotor areas, as well as within temporal areas, and between medial and visual areas (Fig. 4 C). In the distance metric calculated from the result, several compact regions with low pairwise distance emerged (Fig. 4 D).

The result (Fig. 4E) led to eight subregions, where four of them covered the majority of the prefrontal, medial, anterolateral and visual modules of Harris et al. (2019) respectively. However, they also contained significant parts of the temporal module, and the somatosensory and motor areas. Consequently, the remaining four subregions covered only a small “core” of the MOs, primary somatomotor, SSs and temporal regions respectively. More formally, we calculated the intersection-over-union of the established parcellation indicated in Fig. 4E with the regions resulting from the split at reversals (Fig. 5B). We used the result to assign tentative names to the regions (Fig. 5C).

**Figure 5:**
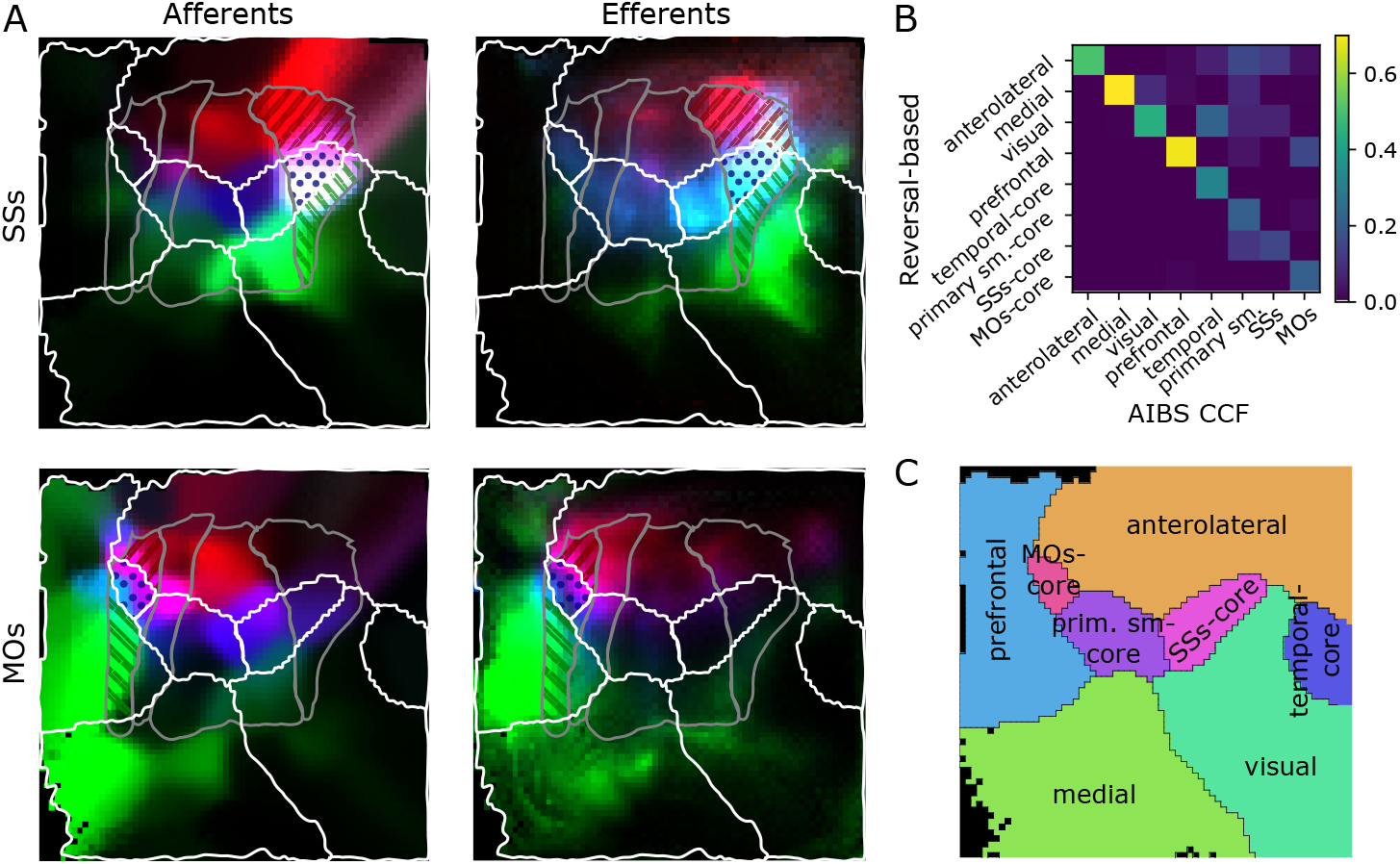
Evaluating the first split against the established parcellation. A: Locations of afferent (left) and efferent (right) cortical locations of regions in the established parcellation, when intersected with our parcellation. Grey lines: Established split into primary and secondary somatosensory and motor regions. White lines: Proposed parcellation of our first split. Top: Region “SSs” is split into an anterior (red stripes) core (blue dots) and posterior (green stripes) part by our parcellation. Afferent / efferent locations are indicated in red, blue and green respectively. Bottom: “MOs” is similarly split into three parts. B: Intersection over union of regions resulting from our first, reversal-based split (vertical) and the established parcellation as indicated in Fig. 4E. C: Tentative labels for the resulting regions.

Curiously, the primary somatomotor regions were split up between six of the eight subregions detected from reversals (Fig. 4E). The apparent lack of such a fundamental level of organization as clearly delineated somatomotor regions led us to investigate more closely.

First, we found that the split of somatosensory areas between five reversal-based regions closely mirrors the split into regions associated with body parts of the established parcellation (Fig. S7A vs. B). The intersections with our anterolateral, “SSp-core” and “SSs-core” regions correspond to the mouth-, nose-, and upper limb-related parts respectively. The intersection with our visual regions corresponds to the barrel field, and medial regions to a combination of lower-limb and trunk-related parts. We conclude that reversal detection recreates the internal parcellation of somatosensory regions, but not its external borders.

Another notable result was the split of the MOs and SSs regions of the established parcellation into three subregions, located anterior, central, and posterior, respectively. We investigated the incoming and outgoing connectivities of these subregions, considering whether there is a fundamental difference in their organization, or whether they are better described as single homogeneous regions (Fig. 5A).

Unsurprisingly, each subregion interacted more strongly with its direct neighbors, but there were also fundamental differences in their long-range connectivity. While the anterior part of SSs receives its strongest inputs from anterior neighbors, the posterior part samples from medial neighbors (specifically, from the barrel field in the established parcellation). While the posterior part of MOs sends and receives extensive connectivity from prefrontal and medial areas, largely avoiding somatosensory regions, the anterior part interacts mostly with the anterior parts of primary motor and somatosensory regions. The core parts of both regions interacted mostly with each other and the core parts of primary somatosensory regions; this was mostly visible in the efferents of SSs.

Finally, the split of motor regions into subregions is arguably consistent with functional data: Guo et al. (2014) described a role of specifically the anterolateral motor areas (ALM) in an object location discrimination task that is not shared by the remaining parts of the motor regions. The ALM region aligns closely with the intersection of motor areas and the reversal-based anterolateral region.

### 2.5 Building a hierarchy of mouse cortical regions

We continued the applications of the algorithm to detect successive splits and thereby build a hierarchy of regions. We performed a second application of *reversal detection*, followed by three applications of *cosine distance clustering*. As before, we switched methods when *reversal detection* no longer resolved additional granularity. In the following paragraphs we will describe the result and the reasons to stop after five applications of the algorithm.

The resulting hierarchy has 41 leaf regions at six levels (Fig. 6A1). Leaf regions exist at each hierarchy level except for the root, indicating that these regions could not be split further by the algorithms. The sizes of leaf regions vary considerably with the largest one being approximately eight times larger than the smallest (Fig. 6A2).

**Figure 6:**
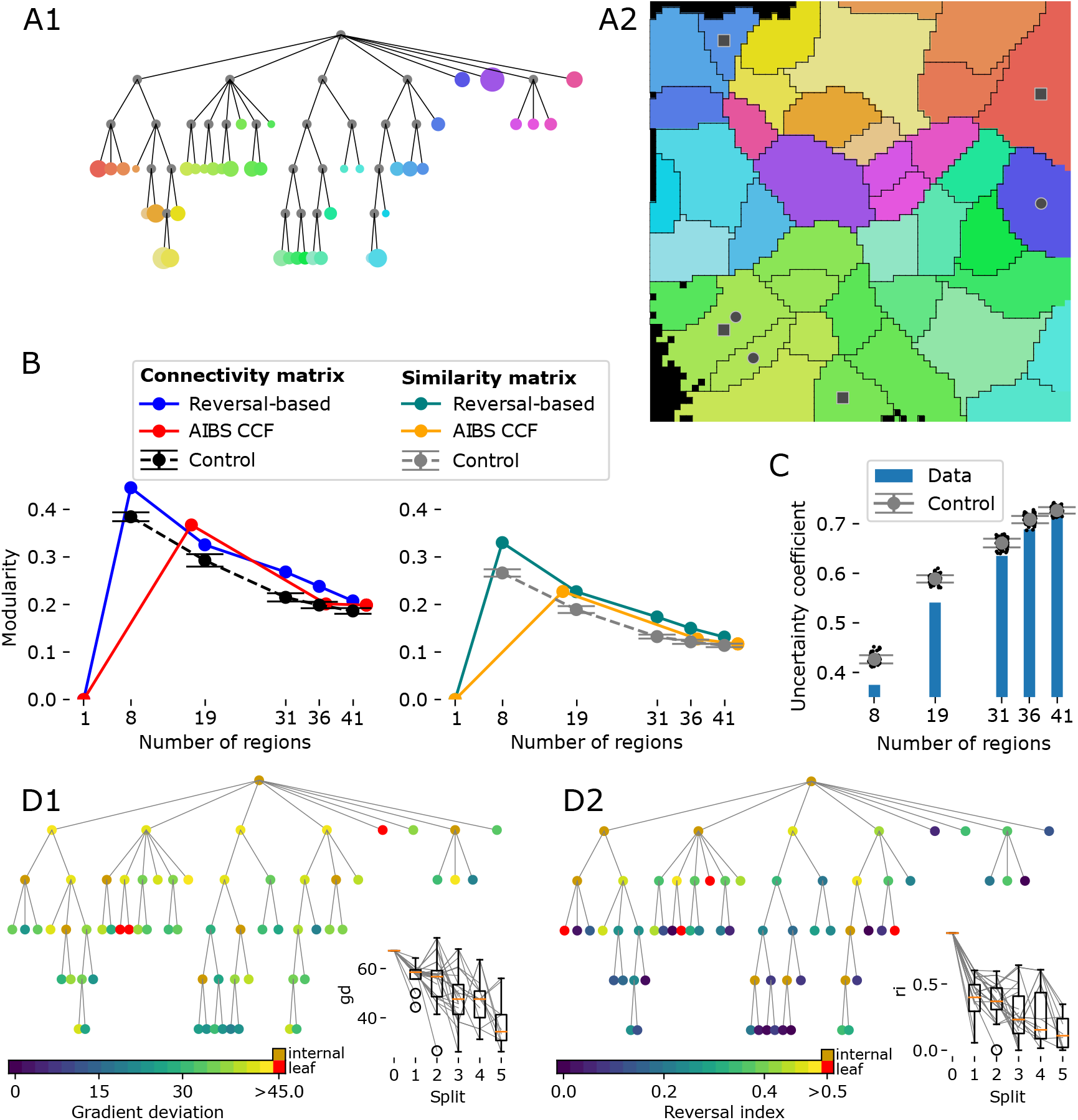
Complete parcellation results. A1: Tree plot of the resulting hierarchy after five successive splits (top to bottom). Size of leaf nodes proportional to region size. A2: Locations of leaf regions. Colors as in A1. B: Modularity of parcellations evaluated with the voxelized connectivity matrix (left) or matrix of similarity of connectivity (right). Plotted against the number of regions at each hierarchy level for the reversal-based parcellation (blue / green), a random parcellation (see Methods) with the same number of regions (black / grey), and the AIBS CCF parcellation (red / yellow). C: Similarity of the AIBS CCF parcellation and the reversal-based parcellation, measured as their uncertainty coefficient (see Methods). Blue bars: values at various hierarchy levels. Black dots and grey error bars: For a random parcellation with the same number of regions instead. D: Values of *gd* (D1) and *ri* (D2) indicated by the color of nodes in tree plots of the hierarchy. Red indicates values above the acceptance threshold, orange if it is an internal node. Insets: Box plots of the values after each split and the changes caused by individual splits (grey lines).

We calculated at each hierarchy level the modularity of the parcellation (see Methods) with respect to the raw connectivity strengths and compared them to corresponding values for the established AIBS CCF hierarchy and parcellation (Fig. 5B, left). As modularity naturally decreases with increasing granularity, we plotted the results against the number of regions and compared them also to 100 random, but spatially continuous parcellations (see Methods; Fig. 5B, black line and error bars). For both reversal-based and established schemes the modularity is significantly larger than in the random controls, but gets closer to them in the lower hierarchy levels. When we calculated modularity based on the (cosine-)similarity of connectivity instead, results were comparable (Fig. 6B, right).

Note that neither scheme aims to maximize the measure. The AIBS scheme unifies multiple historical schemes based on anatomy and neuron function, and our scheme allows very different connectivity within a subregion as long as it varies along a continuous gradient. Yet, this measure shows that our parcellation provides a qualitatively similar solution. This was also evident in a visual comparison of the connectivity matrices (Fig. S7C). Further note that to ensure comparability, we calculated modularity in the flattened view also for the AIBS parcellation; values may differ if calculated between the original voxels.

The demonstrated qualitative similarity between the reversal-based and the established solutions led to the question to what degree they actually describe the same parcellation. To quantify this, we calculated their similarity in terms of the *uncertainty coefficient* (Fig. 6C, for details see Methods). This information theoretical measure yields values between 0 and 1, with 1 indicating a parcellation identical to the established one and values close to 0 a parcellation orthogonal to it, such as the split into cortical layers. As any split into subregions is likely to produce values larger than 0, we compared the results to the same random parcellations as before. We found that values for the reversal-based parcellation were significantly *lower* than for the controls, i.e. it is actually more orthogonal to the established scheme than expected by chance. This indicates that differences between the schemes are due to the reversal method detecting previously unaccounted trends in connectivity and not merely generating random results or picking up noise. However, the trend weakened with number of regions until after five splits the reversal-based solution was within a single standard deviation of the controls with respect to this measurement.

As a final test, we evaluated how the reversal-based parcellation evolves with each split with respect to the previously defined quality metrics (Fig. 6 D). We found a large spread of values for both metrics, but the overall mean and median decreases reliably with each split. Occasionally, a child region results in a larger value for a metric than its parent, indicating a worse solution, but this is typically followed by an even larger decrease in the next split (Fig. 6D, inset, grey lines). Yet, after five splits, three regions were above the defined threshold for *gd* (Fig. 6D1, red; A2, circles), and four for ri (Fig. 6D2, red; A2, squares). All of them appeared before the last split, indicating that the algorithm was not capable of splitting them further, and that additional applications would not lead to an improvement.

We decided to stop after five applications of the algorithm for a combination of reasons: First, at that point the parcellation was close to a random parcellation in terms of modularity and uncertainty coefficient (Fig. 6B, C). Second, the number of new regions resulting from the split had stagnated with most regions already being “atomic”. Third, remaining regions with values for *gd* and *ri* exceeding the threshold could not be split by the algorithm.

## 3 Discussion

We have demonstrated that gradient reversals at the borders between adjacent sensory regions can also be found in the connectivity of voxels belonging to the regions. We have further demonstrated that these gradients can be isolated through the diffusion embedding technique. This approach differs conceptually from previous approaches based on clustering of matrices of connectivity similarity. Clustering-based approaches are based on the idea that connectivity profiles of voxels within a region are homogeneous or at least similar. However, as pointed out by Eickhoff et al. (2015), functional gradients often exist within regions, as shown in earlier studies (Wandell et al.,2007; Hensel et al., 2015). As they are likely reflected by corresponding connectivity gradients, this contradicts the underlying assumption of similarity clustering. Conversely, our approach allows for dissimilar profiles, as long as the dissimilarity is the result of a continuous gradient within the region, reversing at its border.

We have developed an algorithmic pipeline based on this approach that largely automatically breaks up a voxelized connectivity structure into a partition of the volume into subregions (reversal-based parcellation). In addition to the partition, the solution also provides a tentative hierarchy of subregions, although evaluations on a test model show that individual levels of the hierarchy may remain incomplete. At the same time, the evaluations demonstrated a very large degree of robustness against unstructured noise in the input connectivity data. We also developed quality metrics that can be used to detect algorithm failure. They can also be applied to other parcellations to evaluate how close they come to an ideal reversal-based solution.

At the core of the pipeline is the automatic placement of region border, which we implemented in with two different approaches. Of these, *reversal detection* was more conservative, requiring more successive applications and not resolving the full granularity (Fig. 3B1; S4C2). This was because it requires a sub-region to be completely encircled by reversals to be detected. Conversely, *cosine distance clustering* was more affected by noise, especially when tasked to split large sub-volumes (Fig. 3B2). Therefore, we found that the best approach combines the two, beginning with *reversal detection* until no further granularity can be resolved, followed by *cosine distance clustering*. Both approaches are limited in terms of spatial resolution by the numerical calculation of a spatial gradient. This makes it harder to resolve narrow and elongated shaped regions. Regions narrower than three times the voxel resolution are theoretically impossible to resolve, and in practice the limit is even higher.

Beyond these algorithmic issues, one potential danger lies in the hierarchical nature of the algorithm combined with the context-dependence of the detected gradients (Sec. 2.2). This means that great importance is given to first few splits of the target volume, as they determine the context used for all future splits. Should the algorithms fail in the early steps, the entire parcellation would be affected. Other algorithms fail less drastically, with errors at one location not necessarily affecting others.

Finally, one limitation of the presented approach is that it requires a way to project voxel positions into the plane, with the resulting parcellation always being orthogonal to the projection. This does not readily generalize to all brain structures. Mathematically, all techniques we employed could be generalized to work in three dimensions instead, allowing their use in regions where a flattening is harder to define. *Cosine distance clustering* can be expected to work in three dimensions as well, while it remains unclear whether detected reversals would encircle subregions also in three dimensions.

We applied the algorithms to the voxelized connectivity of mouse isocortex (Oh et al., 2014; Knox et al., 2019). The resulting parcellation differed from established schemes, but did so in a demonstrably nonrandom way. Further, when evaluated in terms of modularity, it was further away from a random solution than the established scheme. This indicates that the reversal-based parcellation captures novel connectivity trends that cannot be detected when connectivity is evaluated with respect to the established parcellation. One example is a specialization of the anterolateral motor regions that was not present in the established parcellation, but had been described in earlier literature (Guo et al., 2014). This demonstrates the importance of also considering a parcellation based on connectivity.

The parcellation presented is not to be understood as a replacement of existing schemes, but as complementary. Which one is superior depends on the use case. For example, approaches to connectivity-based parcellation that maximize modularity will by design yield sub-regions with mostly internal connectivity. This is useful for identifying units that can be described in isolation from the rest of the brain, for example in modelling. On the other hand, our scheme is more useful in use cases that require continuous gradients in a region. For example, mouse brain connectivity has been modeled using a single continuous topographical mapping per pair of regions (Reimann et al.,2019). The presence of several reversing gradients in the MOs and SSs regions of the AIBS CCF parcellation led to inaccuracies of the model.

Outside of specific applications, a parcellation is expected to reveal something about the organization of the brain. As it is neuron function that we are trying to understand, an argument can be made in favor of function-based parcellation. On the other hand, connectivity as one of the underlying causes of neuron function (van den Heuvel et al., 2015) puts it closer to the root of the mechanisms we are trying to decipher. Here as well, both approaches can be used in conjunction. For example, one can contrast function-based parcellation, where primary sensory areas are prominently separated with our results, that lack such a delineation. It is important to consider that the presented results were based on intra-cortical connectivity only, while primary sensory areas are largely defined by their thalamic input sources (Sherman, 2016). As such, that aspect of the functional parcellation reflects the quality of bottom-up inputs, and our results are more aligned with the structure of intracortical processing. By considering both, we can improve our understanding of the structure-function relation.

In this context, one advantage of our approach is its adaptability with respect to the types of connectivity to be considered. It would be possible to generate a parcellation based on thalamo-cortical inputs that we predict to feature primary sensory areas prominently. Using corticofugal connectivity and comparing the results may reveal even more aspects (Usrey and Sherman, 2019). Yet, even with only intra-cortical connectivity considered, our result is useful for understanding the organization of mouse intracortical connectivity. This can be in the form of analyzing large-scale *in vivo* recordings of neuronal activity with respect to our parcellation to test which aspects of neuronal function are aligned with it.

The adaptability of the approach extends to the use of other data sources. The set of algorithms can be readily applied to any source of voxelized connectivity in any organism, including undirected connectivity, such as functional connectomes. Indeed, comparable gradients have been found in human functional connectivity data (Katsumi et al., 2021). As functional connectivity can be derived for single individuals, our technique could be used for assessing inter-individual differences, or changes resulting from disease and injury. Derived parcellations come with a distance metric in the form of the uncertainty coefficient, and differences could be easily interpreted in the form of for example growth and shrinkage of corresponding regions. In fact, the algorithms can be easily adapted to data outside of connectivity, working with any voxelized data source where similarity of pairs of voxels can be defined and calculated. This encompasses for example gene expression data, although it is unclear whether this data is organized around spatial gradients with reversals.

Simultaneously a strength and a weakness of the results presented here, is their provenance from a single data source. This ensures a certain level of standardization across the entire cortex. The splitting algorithms also treat each cortical location equally, leading to a result that avoids the potential biases arising when different parts of the parcellation are based on different sources and different techniques. On the other hand, the result will be affected by any bias or weakness in the source data itself. This can be improved by combining several sources of connectivity data or considering both anatomical and functional reversals simultaneously.

## 4 Methods

### 4.1 Diffusion embedding

In short, the method considers a diffusion process along the edges of a graph, described as a Markov chain with a transition matrix determined by the normalized (cosine-) similarity of incoming and outgoing connectivity of each voxel. The embedding coordinates are then given by the eigenvectors of the transition matrix, scaled by their eigenvalues.

Let *C_[i,j]_* be the strength of connectivity between brain voxels *i* and *j*, 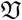 be the set of voxels in the volume to be partitioned, 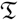 the volume to be considered as a connectivity target, and 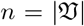. In this work, we considered intra-cortical connectivity exclusively, hence 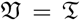, but other options are possible, such as partitioning cortex based on its connectivity with thalamus.

The connectivity profile *P* of each voxel was then given by the concatenation of *C* with its transpose, such that each row of the resulting *n* × 2*n* matrix corresponded to the incoming and outgoing connectivity of a voxel in 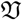.

*P′* was a normalized version of *P* and *S* the cosine similarity of *P′*:

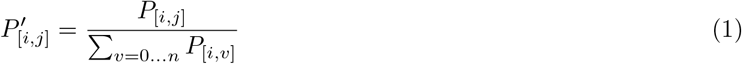

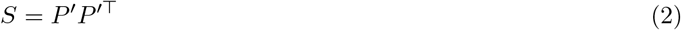

In a final normalization, each entry of *S* was scaled, based on product of the sums of values in the corresponding row and column as follows:

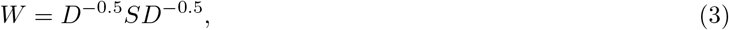

where *D* was a diagonal matrix with each entry *D_[i,i]_* being the sum of the corresponding row of *S*. *W* was then used as the transition matrix, which with this last normalization approximates the Fokker-Planck diffusion (Coifman and Lafon, 2006).

The embedding coordinates used were then the eigenvectors of *W*, scaled by their eigenvalues (*λ^t^*; though we used exclusively *t* = 1). Coordinates were sorted by their eigenvalues and the strongest ones considered as described in the rest of the manuscript. For the steps from equation 3 on we used the implementation of github.com/satra/mapalign, as described in (Coifman and Lafon, 2006; Langs et al., 2015) with parameters *α* = 0.5 and *t* = 1.

### 4.2 Gradient deviation and reversal index

We defined two quality metrics for proposed parcellations based on the gradients found within each of their regions. First, *gradient deviation, gd* evaluates the orthogonality of the two strongest gradients in a region *κ*:

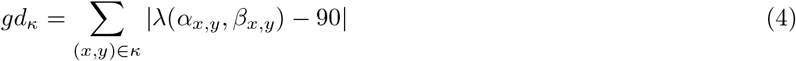

where λ(.,.) measures the angle between two gradients, here the two strongest gradients *α*, *β* at pixel (*x, y*).

The second metric, *reversal index* evaluates the continuity of the gradients, i.e. the absence of reversals. It first counts the pairs of pixels with an angle above 90 degrees in the same component *ω:*

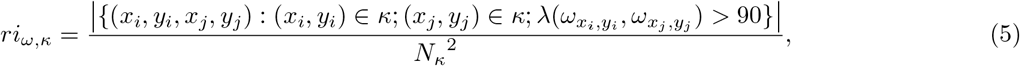

where *N_κ_* refers to the number of pixels in *κ*. Then, this measurement is summed over the two strongest components:

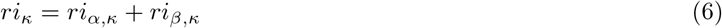

### 4.3 Algorithmic detection of reversals

Two approaches automatically detected gradient reversals and drew borders between regions separated by them: *cosine distance clustering* and *reversal detection*.

Both begin by applying diffusion embedding (Vos de Wael et al., 2020) to the matrix of directed connection strengths between voxels, extracting the 20 strongest components (Fig. 2A, left). As discussed in the introduction and Fig. 1, we work in a flattened view of the volume of interest; as such we average the value of each component over the voxels mapped to the same pixel. We then calculate their gradients with respect to the grid of pixels. The next steps differ between the two approaches.

*Cosine distance clustering* (Fig. 2A, bottom) considers the gradients of the strongest twenty components and directly calculated a distance metric based on them. For each pair of pixels, a weighted sum of cosine similarities of the gradients was considered, where the weight was equal to the strength of the associated component. First the gradient of the nth component (*ω_n_*) at a location i was normalized (as in Fig. 1F:

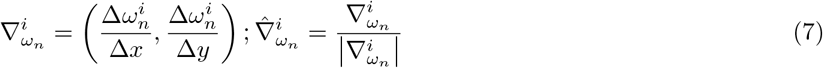

Then the weighted sum of cosine similarities was calculated:

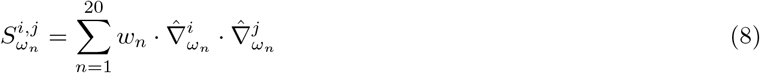

This was then converted to a distance by subtracting the value from the maximum possible value

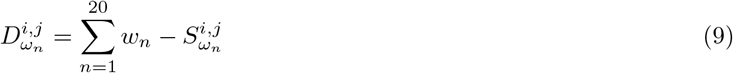

We then applied the HDBSCAN algorithm (Campello et al., 2013), resulting in a hierarchical clustering that yields an initial parcellation.

*Reversal detection* (Fig. 2A, bottom) considered the normalized gradient of a single component in its flattened view, 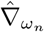, as above. We calculated the degree of reversal at each location by convolving the gradient field with a two-dimensional Gaussian kernel. If gradients around a pixel have the same orientation, the vector resulting from the convolution at that point will also have unit length; conversely, a reversal would lead to gradients with inverted orientation cancelling each other out in the convolution. We considered a convolved vector shorter than 0.97 to indicate that the location was part of a border. We then determined contiguous regions encircled by the same boundary in the following way: We began by building a graph where each node represented a pixel and edges were placed between all pairs of neighboring pixels (direct or diagonal neighbors). The length of an edge was 1 if either of the pixels was tagged as a border, 0 otherwise. We calculated distances between all pixels as the length of the shortest path in the graph between the pair, as calculated by the Dijkstra algorithm (Dijkstra and others, 1959). Finally, we generated groups of pixels by running the Ward linkage algorithm on the matrix of pairwise distances. We chose the number of clusters between 2 and 10 by optimizing the resulting silhouette score.

Unfortunately, reversals might not be observed equally on every diffusion component, so a first operation is to determine which component to use to perform the reversal detection. While an intuitive choice may be the first, i.e. strongest, component, in practice we found that: 1) This component was not guaranteed to yield the most useful split; and 2) the following components often had very similar strengths, indicating that their order may be partially arbitrary. Consequently, we decided to run the algorithm on all of the 20 strongest components and choose the one that minimizes the *reversal index*, *ri* within the resulting subregions:

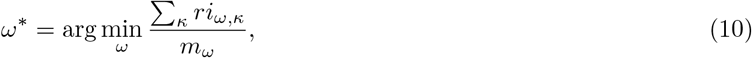

where *m_ω_* is the number of subregions resulting when a split is applied based on *ω* and *ri* is defined as in equation 5. Note that in actual biological data, the 20th component still had a strength of 10% of the first component.

### 4.4 Post-processing the initial split

The shortcomings of both methods required a number of post-processing steps (Fig. 2A, right). First, unlabelled pixels (see above) needed to be classified. To that end, we train a Support Vector Machine to classify the unlabelled pixels based on their *x* and *y* coordinates in the flat coordinate system (Fig. S6A). This step serves not only to extrapolate the first classification, but also to generalize the output to a less noisy result. Indeed, due to potential noise in the input data, the resulting parcellation of the first gradient classification can be irregular and scattered, especially for *cosine distance clustering*. Second, at this stage all pixels are classified but some clusters of the same class can be spatially non-continuous, therefore these clusters must be differentiated. This is done by hierarchical clustering of pixels of the same class based on their Euclidean distance (Fig. S6B). Third, we“unflatten” the resulting parcellation from pixels to voxels by looking up the reverse images of each pixel (Fig. S6C). Finally, we specify a minimum size threshold for the resulting regions to reject any regions that are unrealistically small and merge them with their closest neighbour (Fig. S6D).

### 4.5 Building toy models of connectivity

We built models of connectivity inside a voxelized 3-d cube, with a regular grid of a given resolution and depth specifying a spatially nested hierarchy of regions. Each level of the hierarchy split the cube along one axis, alternating between the y- and z-axis (Fig. 2B). Connection strengths were generated for a hierarchy level by reversing the diffusion process. We prescribed continuous and reversing target values of the two strongest components, *α* and *β*, i.e. attributing increasing – or decreasing – component values to voxels and reversing these components where a different region starts (Fig. 2B, right). E.g. for a split along the y-axis:

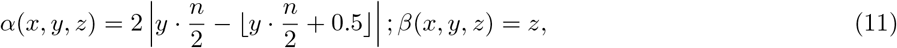

where *n* is the *resolution* of the model, i.e. the number of subregions in the hierarchy level.

The connection strength between each pair of voxels *i* and *j* was then defined based on Euclidean distance of those values:

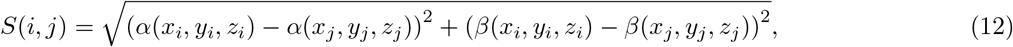

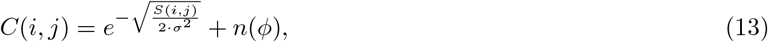

where *σ* was set to 0.1 times the maximum of *S*(*x, y*) and *n*(*ϕ*) denotes noise with a uniform distribution between –*ϕ* and *ϕ* that was added to test the resistance against noise.

This process could be applied at a given level of the hierarchy to connect the subvolumes at that level of the hierarchy (Fig. 2B, right). This yielded one connection matrix per hierarchy level. We combined matrices for all hierarchy levels by piecewise multiplication of their entries (*reversing hierarchy model*, Fig. 2C1). As an alternative approach, we only considered the connection matrix for the lowest hierarchy level. Then, for each entry we considered the pair of regions it corresponds to, looked up their path distance in the hierarchy tree, and divided the value of the entry by that distance (*node distance model*, Fig. 2C2).

We built more complex, random models (*Random split model*) based on recursively splitting the volume at randomly drawn lines. For each split, first the angle of the line is randomly drawn. It is drawn with uniform probability from [0, 2*π*] for the first split and from 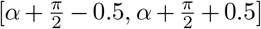 for all following, where *α* is the angle of the previous split. The location of the line is then randomly determined such that the sub-volume is split between 40%-60% and 60%-40%. The process is repeated for each resulting sub-volume until they are below a size threshold, or until either a specified number of regions has been generated.

Connectivity was based on the location of each placed line. Each line defined a mirror operation. Points mirrored on top of each other are then set to project to each other in the following way: Let P, P′ be a pair of points mirrored on top of each other. The strength of connections from P is then given by:

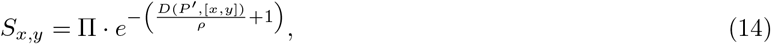

where Π and *ρ* depended on the hierarchy level *L* of the split, beginning at 1 for the first split.

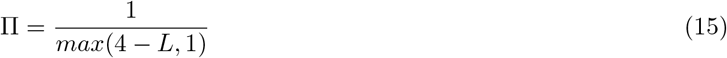

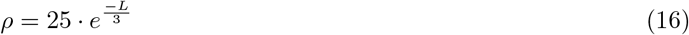

### 4.6 Calculation of the uncertainty coefficient of pairs of parcellations

A parcellation *X* associates each voxel *v* with a region 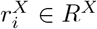. We describe this association as *X*(*v*) = *i*. Let 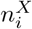 be the number of voxels associated with 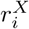 and 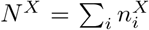 the total number of voxels. We can then consider *P^X^*, the probability distribution given by the region associated with voxel that has been randomly picked (with uniform probabilities):

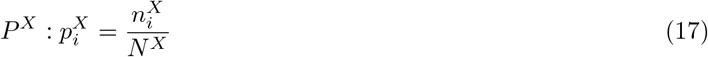

As with any probability distribution, we can calculate its entropy:

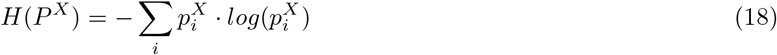

Given two parcellations *X, Y* that are defined on the same set of voxels, we can similarly consider their joint distribution *P^(X,Y)^* where 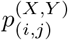 is the probability that a random voxel is assigned to region *i* in *X* and to *j* in *Y*.

This allows us to calculate the mutual information of the probability distributions associated with a pair of parcellations. This is the reduction of entropy of one of them when the value of the other is revealed. That is, if you know the region identity of the random voxel according to one parcellation, how much does it help you to guess its identity in the other?

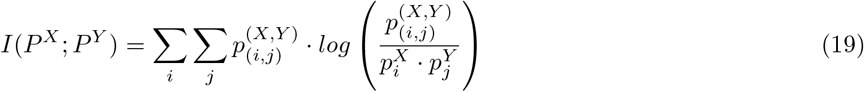

Normalizing it by the entropy of *X* yields the *uncertainty coefficient*:

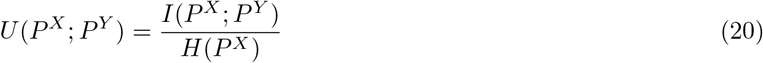

This is a measure of similarity between parcellations. Intuitively, if revealing that the identity of a voxel according to *Y* is *j* yields a lot of information about its its identity in *X*, this means there must be at least one region in *X* that overlaps strongly with *j*. Mathematically, we can see how for *X* = *Y* equation 19 equals equation 18 because 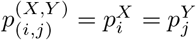 for *i* = *j* and 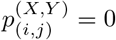 for *i* = *j*. Consequently, in that case *U*(*P^X^; P^Y^*) = 1.

Conversely, if *P^X^* and *P^Y^* are statistically independent, then by definition 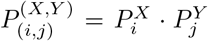 and consequently *I*(*P^X^; P^Y^*) = *U*(*P^X^; P^Y^*) = 0. This is the case for orthogonal parcellations.

In the manuscript we set *X* to the established AIBS CCF parcellation and Y to whichever parcellation or control parcellation we want to evaluate.

### 4.7 Calculation of the modularity of a parcellation

The modularity of a parcellation with respect to a matrix of connection strengths is the difference between the fraction of connection strength that is found between pairs of voxels in the same region and the fraction of the number of pairs of voxels in the same region.

Let *X* be a parcellation as defined in the previous section. Let *M* be the connectivity matrix, such that *M*(*v, w*) yields the strength of connectivity between voxels *v* and *w*. Then the fraction of connection strength found within a region is:

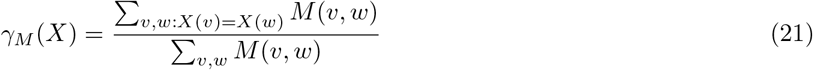

The fraction of voxel pairs that are in the same region, relative to all possible pairs is:

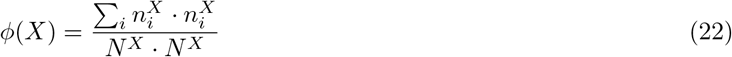

The modularity is then the difference between the two, _*γM*_(*X*) – *ϕ*(*X*). As individually their values are between 0 and 1. This measurement is between −1 and 1.

If connectivity strengths are independent of the parcellation, then the expected value of the measurement is 0, if strong connectivity is found predominately within a region, the measurement is > 0.

### 4.8 Generation of random control parcellations

We generated comparable, but random parcellation schemes the following way: First, a random control is paired with the reversal-based parcellation after a given number of splits in order to match its number of resulting subregions. For a given number of subregions, we generated 100 random parcellations with two different algorithms (50 per algorithm).

First, we generated 50 schemes by first randomly distributing a number of points equal to the target number of subregions within the axis aligned bounding box of the pixel positions of the flat view. Points were randomly picked with uniform density within the bounding box. Then, pixels were classified into subregions based on the random point they were closest to. Second, we generated 50 schemes according to the *random split model* described above in Sec. 4.5. Both controls generated random parcellations with spatially continuous subregions of various sizes and the same number of subregions as the reversal-based parcellation.

## Data and code availability

The resulting parcellation and hierarchy are available under the following DOI: 10.5281/zenodo.7032168

The code used to create the parcellation from brain connectivity data is available at: github.com/MWolfR/ConnecMap

The connectivity data used was the voxelized mouse connectome published by the Allen Institute for Brain Science, available at connectivity.brain-map.org (data) and github.com/AllenInstitute (code).

## Acknowledgements

This study was supported by funding to the Blue Brain Project, a research center of the Ecole polytechnique fédérale de Lausanne (EPFL), from the Swiss government’s ETH Board of the Swiss Federal Institutes of Technology.

The authors would like to thank Daniela Egas-Santander for helpful suggestions and discussions.

## Supplementary Material

**Figure S1:**
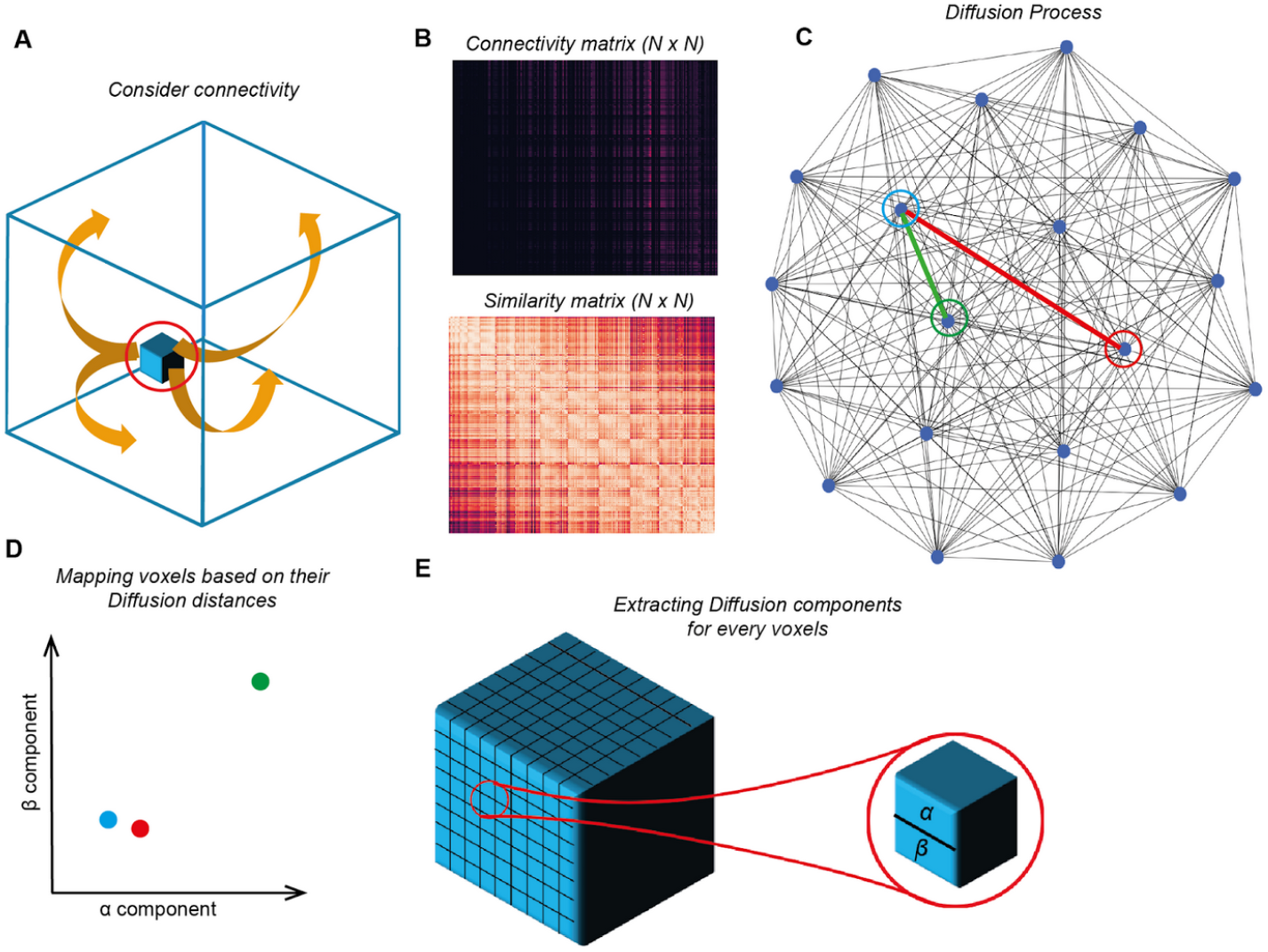
Framework of the diffusion embedding. From a connectivity matrix between source voxels and target voxels, we build a similarity matrix based on how the voxels’ connectivity profile is similar (A-B). On this matrix, we run the diffusion process which will strengthen high connected pathways (red) and weaken connected pathways (green) (C). This process allows to reveal an embedded geometrical space of connectivity, where strongly connected voxels are close to each and in opposite poorly connected voxels are far from each other (D). To flatten this geometrical space, we therefore extract the 2 dimensions associated with the 2 strongest eigenvectors, which we call alpha and beta components (E).

**Figure S2:**
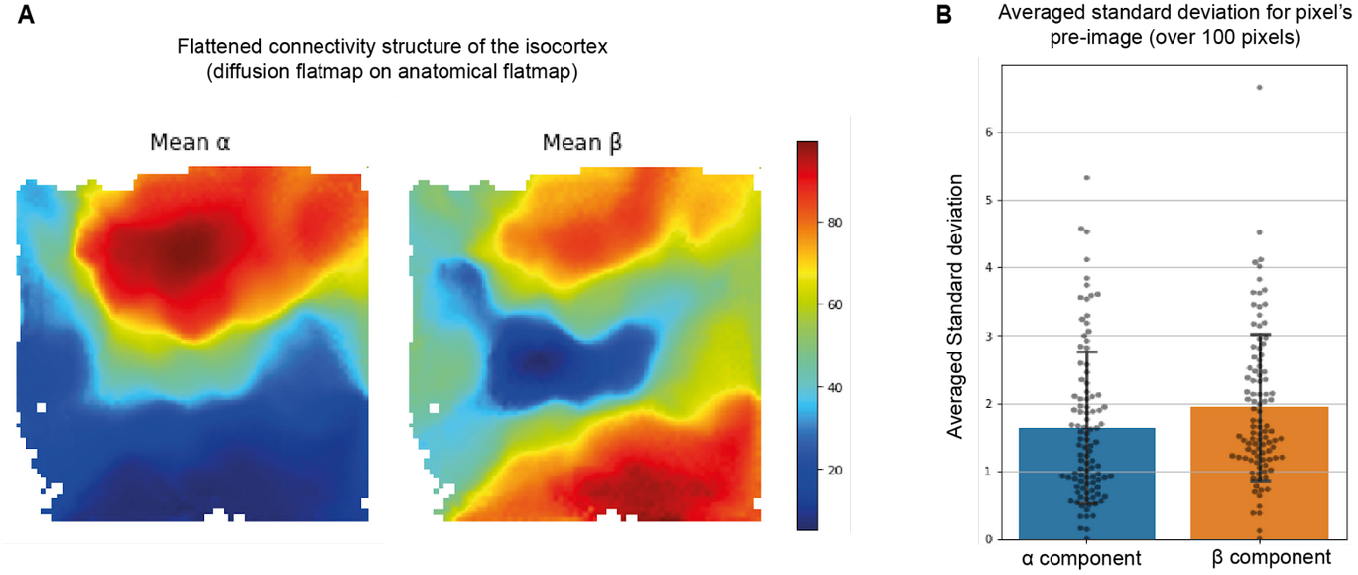
Projecting values of connectivity components to two dimensions. (A) Flattened reconstruction of the connectivity structure of the mouse isocortex by averaging the diffusion components of voxels corresponding to a pixel. (B) Standard deviation of the diffusion components of voxels corresponding to a pixel, averaged over 100 pixels.

**Figure S3:**
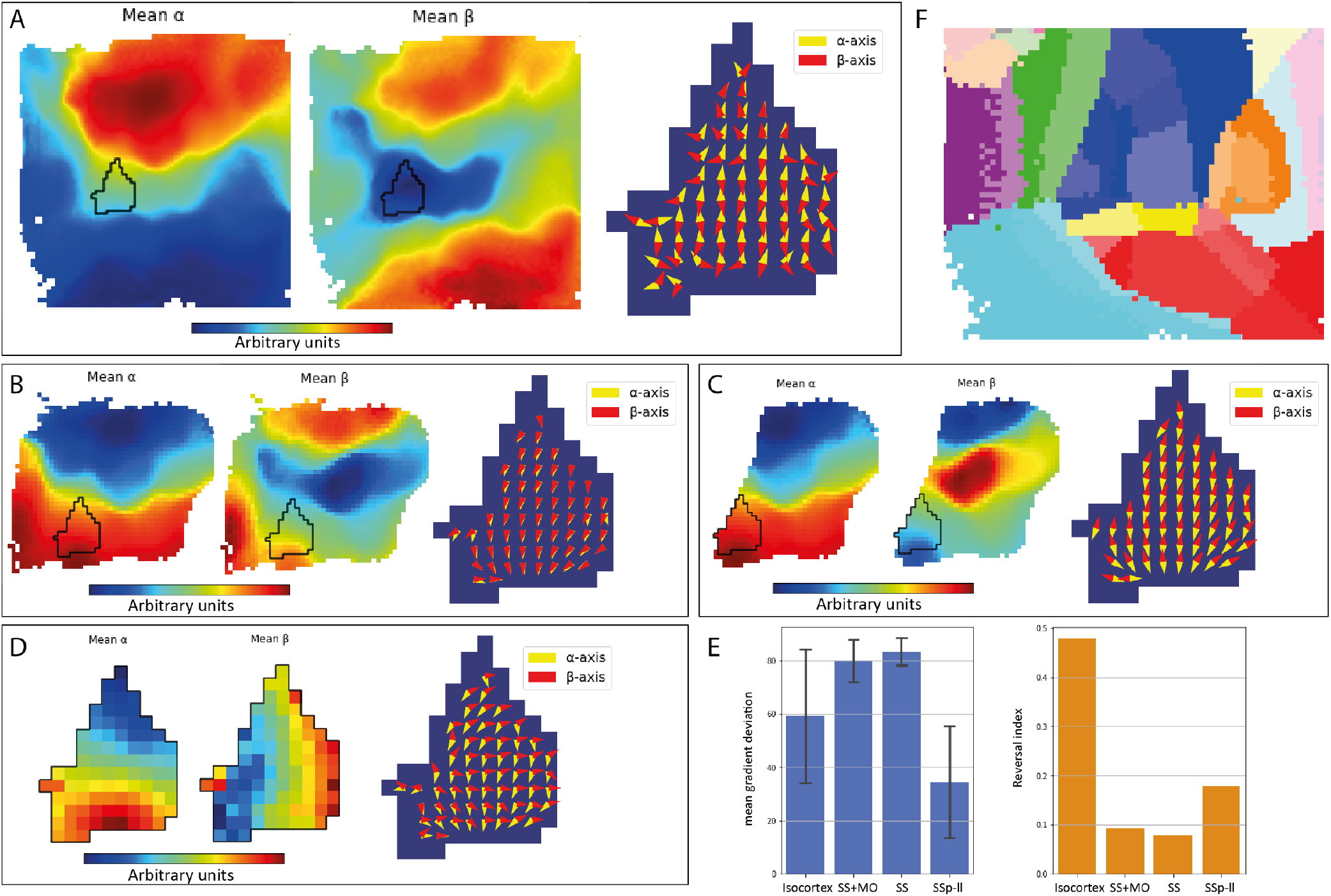
Diffusion embedding with different spatial contexts. A: Diffusion embedding performed on the whole isocortex, SSp-ll is annotated by the black line. B: Diffusion embedding of the Somatosensory and Somatomotor areas (blue and green in the parcellation in E.) C: Diffusion embedding of the Somatosensory areas (blue in the parcellation E.) D: Diffusion embedding of the SSp-ll. E: Mean gradient deviation (error bars = standard deviation) and reversal index of SSp-ll at different scales. Even though the connectivity structure of SSp-ll is homogeneous and continuous according to the gradient deviation and reversal index, it can only be revealed by isolating it after successive splits, hence the motivation to split the isocortex several time around gradients’ reversals detected thanks to our method. F: Hierarchical organization of the AIBS CCFv3.

**Figure S4:**
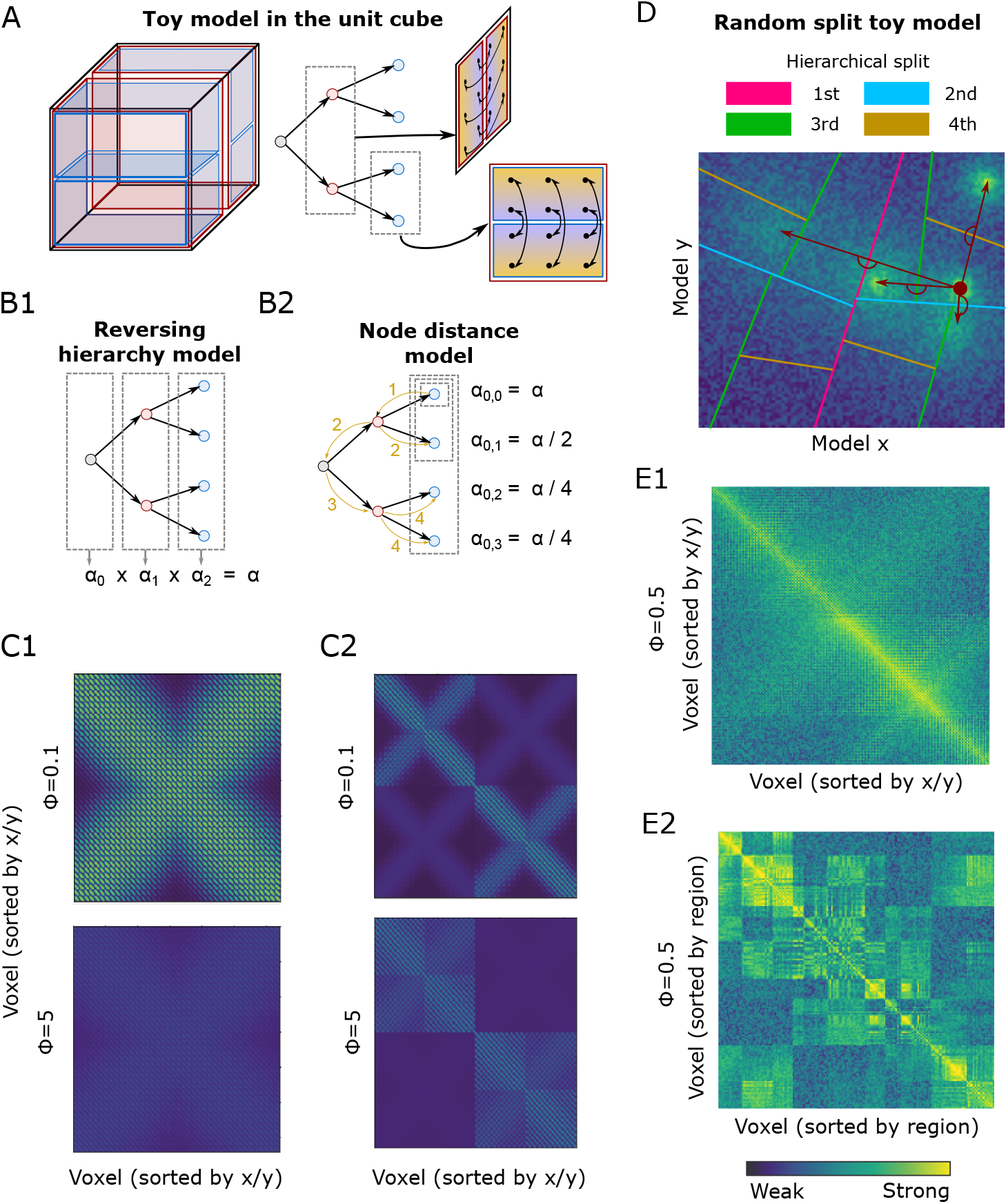
Toy models for the evaluation of the algorithm. A: Left: A model hierarchically splits the unit cube into equal quadrants. Right: The value of the strongest connectivity component is then prescribed as a linear gradient inverting at each border. Connectivity is then based on similarity of the prescribed connectivity components. B: The method yields one connection matrix at each hierarchy level. We construct a merged matrix by either multiplying the matrices at all levels (*reversing hierarchy model*, B1), or by considering the matrix at the lowest level and dividing the values in individual submatrices by the path distance between the nodes they represent in the hierarchy graph (*node distance model*, B2, distances from indicated in yellow). C: Connection matrices of the two models with low (*ϕ* = 0.1) and high (*ϕ* = 5) noise added. D: More complex models are randomly generated by recursively splitting the space in two with lines drawn at random angles. At each split, the points on one side of the line are set to project towards locations at the mirrored opposite side (green arrows for an exemplary source indicated by a red dot). The range around the destination decreases, but strength increases with successive splits. White noise at *ϕ* = 0.5 was added to projection strengths of each voxel pair. E: Connection matrix of the random parcellation in D. Sorted by x, y coordinates (E1) or by region (E2).

**Figure S5:**
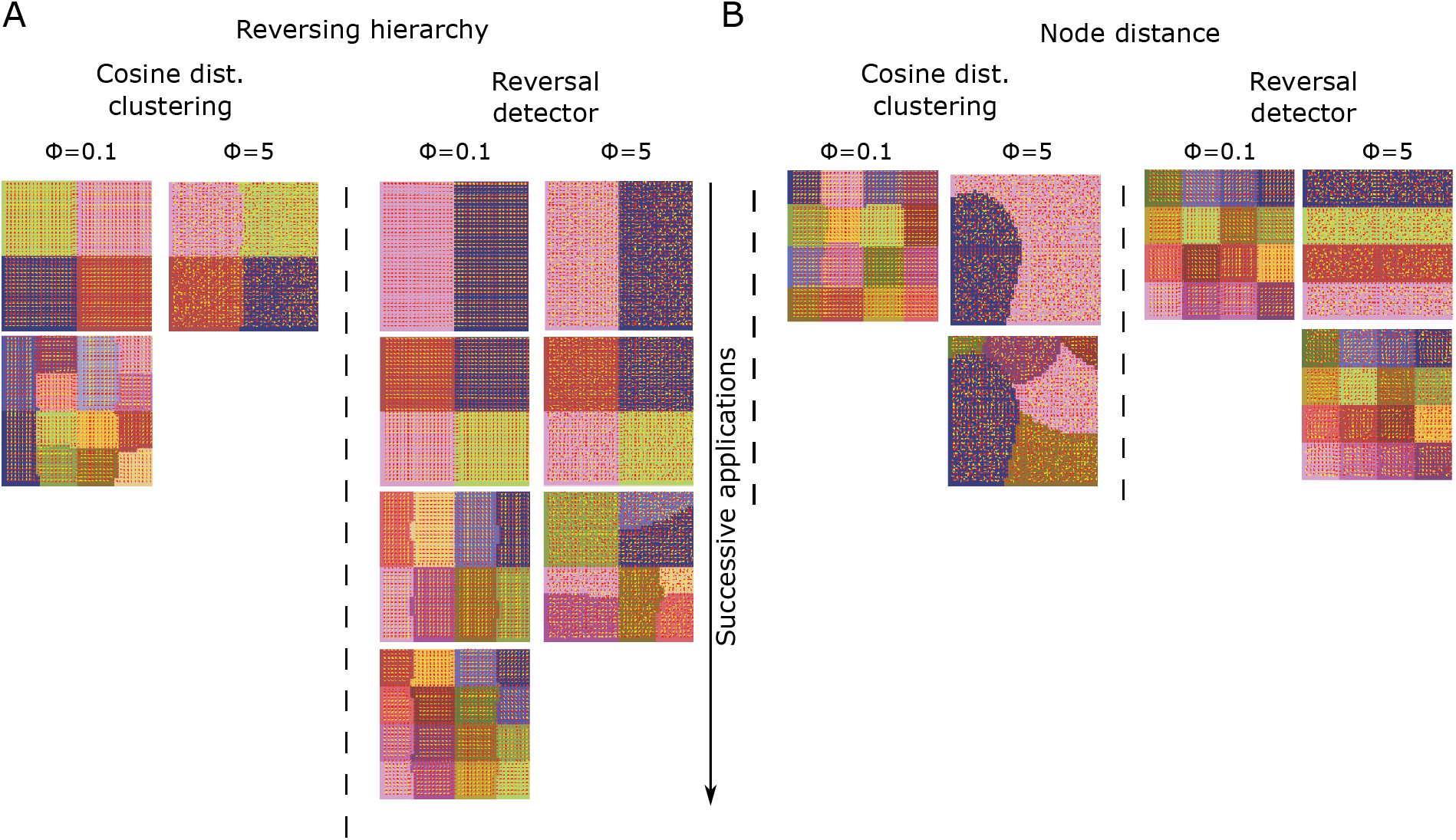
Intermediate results reached while splitting toy models. For the reversing hierarchy (A) and node distance (B) models as indicated in Fig. 3.

**Figure S6:**
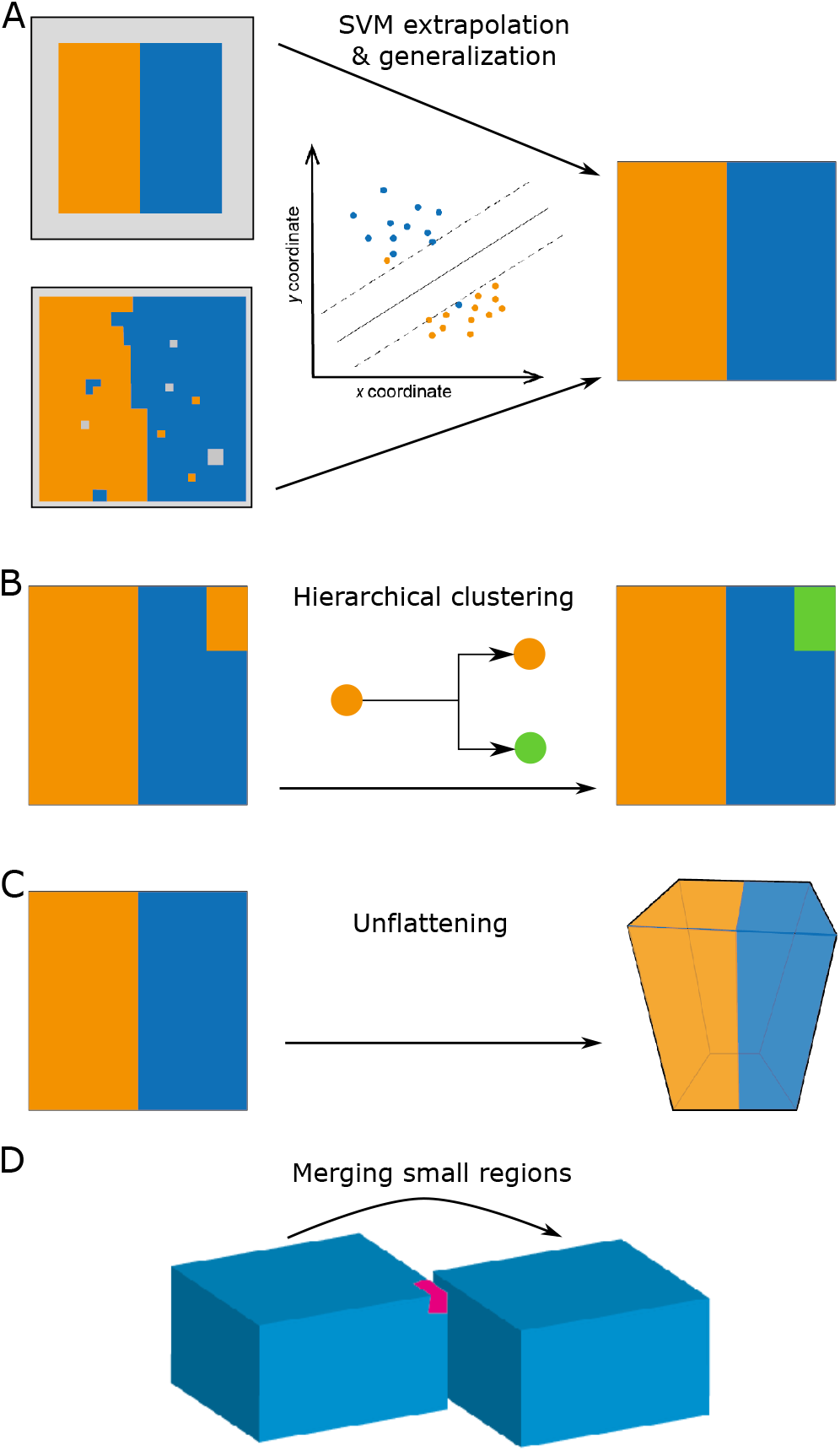
Post-processing of the initial parcellation. Post-processing steps, as indicated in Fig. 2A, that improve the initial parcellation. A: The initial parcellation of reversal detection (top) has many unlabelled pixels (grey) in the periphery. The parcellation of cosine distance clustering (bottom) is noisy and also has unlabelled pixels. A support vector machine is trained that predicts the region label from the location of a pixel. The prediction of the SVM features straighter region boundaries and labels for all pixels. B: Occasionally, the same region label is applied to spatially non-continuous patches. For a given label, hierarchical clustering is applied to the pairwise distances of the location of associated pixels. This splits up non-continuous regions. C: The parcellation is unflattened, i.e. projected back into 3d space by looking up all voxels that are associated with a given pixel. D: Regions below a volume threshold are merged with their nearest neighbor.

**Figure S7:**
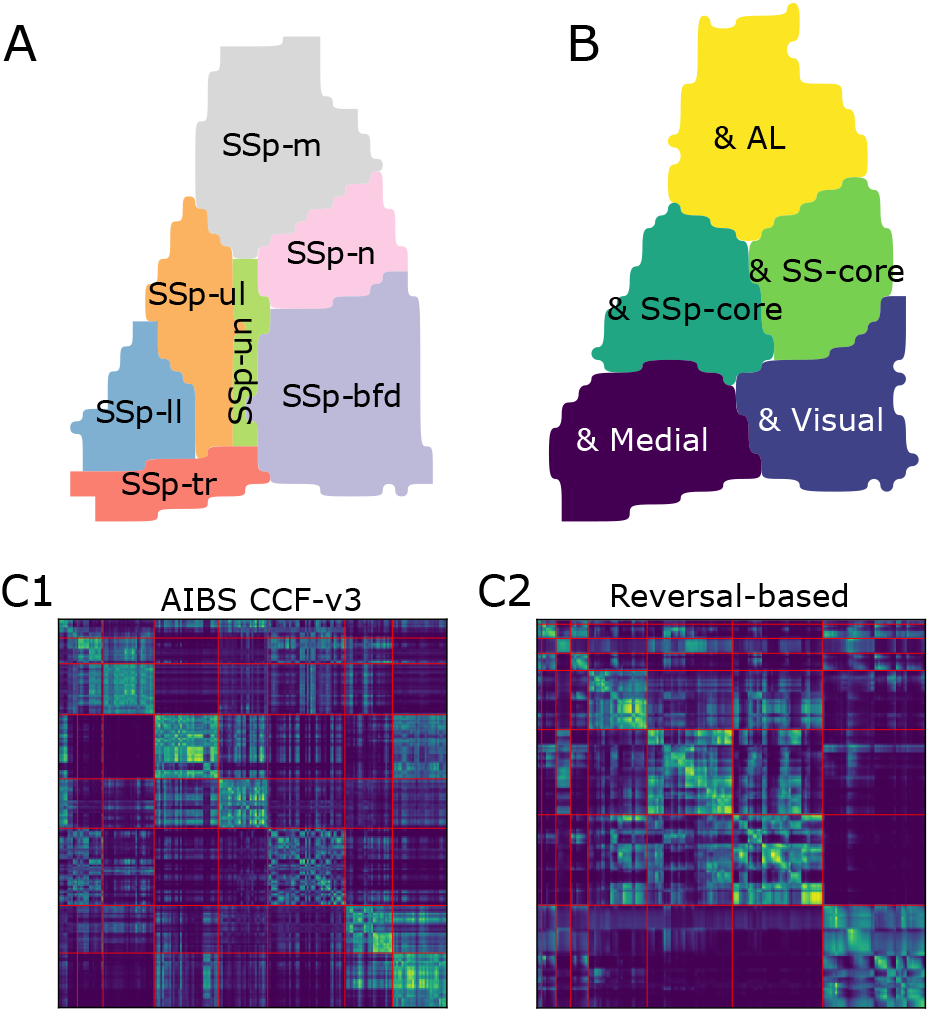
Comparison of the parcellation of somatosensory regions. A: Parcellation of primary so-matatosensory regions, as defined in the established AIBS CCF. B: Intersections of primary somatosensory regions in AIBS CCF with regions at the first hierarchy level of our reversal-based parcellation. C: Matrix of connections strengths between cortical locations. C1: Rows and columns sorted by high-level regions of the established parcellation (as shown in Fig. 4E). C2: Sorted by regions in the first hierarchy level of the reversal-based parcellation. Red lines indicate breaks between regions.

## Notes

https://doi.org/10.5281/zenodo.7032168

### Competing Interest Statement

The authors have declared no competing interest.

### Summary of Updates

Revised to address peer reviewers' comments. This is the final version that will be published soon.

